# Cholinergic impairment in the dorsal motor nucleus of the vagus during experimental Alzheimer’s disease

**DOI:** 10.64898/2026.09.19.752906

**Authors:** Aidan Falvey, Santhoshi P. Palandira, Saher Chaudhry, Joshua J. Strohl, Patricio T. Huerta, Tea Tsaava, Aisling Tynan, Cristina d’Abramo, Luca Giliberto, Jeremy Koppel, Yousef Al-Abed, Michael Brines, Philippe Marambaud, Sangeeta S. Chavan, Eric H. Chang, Kevin J. Tracey, Valentin A. Pavlov

## Abstract

Cholinergic neurons in the dorsal motor nucleus of the vagus (DMN) in the brainstem are a key source of efferent vagus nerve fibers that regulate vital functions, including heart rate and inflammation. Whether the integrity of DMN cholinergic neurons is affected during Alzheimer’s disease (AD) remains unknown. Here, in female and male mice with experimental AD (5xFAD), which exhibit age-dependent memory impairment, basal forebrain cholinergic neurodegeneration, and microglial alterations, we observe a reduction in cholinergic neuron density in the DMN at 6 and 10 months of age. Furthermore, while an important physiological function of DMN cholinergic signaling, such as suppression of heart rate, is preserved in control mice upon electrical DMN stimulation, the extent of suppression diminishes with age in both female and male 5xFAD mice. In addition, while electrical DMN stimulation lowers pro-inflammatory cytokine levels in control mice subjected to endotoxemia, this anti-inflammatory effect is diminished with age in 5xFAD mice, with females showing earlier dysfunction at 6 months. These results reveal previously unrecognized age-dependent cholinergic deficits in the DMN and disrupted brain–to–periphery vagus nerve circuits in experimental AD. These findings advance our understanding of AD mechanisms and are of interest for the development of conceptually novel therapies.

## Introduction

Alzheimer’s disease (AD) is a debilitating and lethal neurodegenerative disorder and the most common form of dementia, which is projected to affect 1 in 85 people worldwide by 2050 (1–3). AD has a serious deleterious impact on patients, close relatives, and caregivers, and is considered “the greatest challenge for health and social care in the 21^st^ century” (3). Despite considerable research and development efforts, AD remains an enigmatic and incurable disorder, dictating the need for a better understanding of its pathogenesis, which would ultimately inform novel treatments.

The memory impairment during AD pathogenesis has been directly linked to neurodegeneration, particularly affecting cholinergic neurons localized in the basal forebrain (2, 4, 5). Cholinergic drugs, i.e., acetylcholinesterase inhibitors - which suppress acetylcholine biodegradation in the brain – are currently used for the symptomatic treatment of cognitive decline in AD (2, 5, 6). However, the integrity of other clusters of cholinergic neurons in the brain during the disease remains largely unknown. A major localization of cholinergic neurons is the dorsal motor nucleus of the vagus (DMN) within the brainstem medulla oblongata. These neurons project long axons within the efferent vagus nerve, the main nerve of the parasympathetic part of the autonomic nervous system. The vagus innervates the heart and other peripheral organs and regulates the heart rate and other vital physiological functions (7, 8). Discoveries over the last two decades have also revealed that efferent vagus nerve cholinergic signaling regulates pro-inflammatory cytokine responses and inflammation (9, 10). Using optogenetic and electrical stimulation in mice with endotoxemia, we identified the DMN as a major source of vagal cholinergic neurons with anti-inflammatory output (11, 12).

While DMN degeneration has been reported in neurodegenerative disorders such as Parkinson’s disease (13), the anatomical and functional integrity of DMN cholinergic neurons during AD remains largely unstudied. Here, to guide such investigation, we first demonstrate a timeline of previously reported behavioral alterations alongside basal forebrain cholinergic neurodegeneration and microglial changes in female and male 5xFAD mice - a widely used model of AD. Then, in these mice, we reveal previously unrecognized anatomical deterioration of DMN cholinergic neurons compared with age-matched wild-type (control) mice. Furthermore, we show that in 5xFAD mice, the suppressive effects of electrical DMN stimulation on heart rate and peripheral inflammation are compromised in an age-dependent manner. These findings highlight alterations in DMN cholinergic signaling in 5xFAD mice, indicating that brain-to-periphery vagus nerve circuits become dysfunctional during experimental AD. These are novel insights into AD pathology that are of substantial interest for conceptualizing future research and developing new therapeutic approaches.

## Results

### Age-dependent memory deficit in 5xFAD mice is associated with basal forebrain medial septum neurodegeneration and hippocampal microglial activation

To specify the time frame for DMN examination in female and male 5xFAD mice, we first assessed the age dependence of characteristic AD features. Memory deterioration is a key hallmark of AD, and specific hippocampal object-location memory impairment has been documented in patients with AD (2, 3, 14). We examined age-dependent alterations in object-location memory, also known as object-place memory (OPM), in 5xFAD mice. Behavioral experiments with 2-, 6-, and 10- month-old female and male 5xFAD mice, as well as control mice, were performed using a previously established OPM protocol (15). Briefly, mice were first familiarized with the testing apparatus, followed by a sample trial with two identical objects in adjacent quadrants of the box. Then, in the choice trial, one of these objects was moved to a novel location. No significant differences were observed between female 2-month-old 5xFAD and control mice, as shown in **Figure 1 a,b**). However, the OPM ratio in 6-month-old female 5xFAD mice was significantly lower compared with WT mice (**Figure 1 c,d**). Similarly, the OPM ratio was significantly lower in 10- month-old female 5xFAD mice than in WT mice (**Figure 1 e,f).** The same age-dependent behavioral alterations were observed in male 5xFAD mice compared with controls. While there were no significant differences at 2 months (**Figure 1 g,h),** significantly lower OPM ratios were found at 6 months (**Figure 1 i,j)** and 10 months (**Figure 1 k,l).** For each age and sex, we compared the total time mice spent exploring both objects during the choice trial and found no significant differences between WT and 5xFAD mice. These results indicate that differences in the OPM ratio were not attributable to overall group differences in total object exploration time **(Supplementary Figure 1).** The hippocampus, particularly the CA1 area, is involved in OPM (16–18). Cholinergic neurons of the medial septum (MS) in the basal forebrain, via the septo-hippocampal pathway, innervate the hippocampus and play a key role in regulating hippocampal memory processes (19, 20). Examination of MS cholinergic neurons showed that while there were no differences at 2 months, the percentage of choline acetyltransferase (ChAT)-positive neurons in the MS was significantly reduced in female and male 5xFAD mice compared with controls at 6 and 10 months (**Supplementary Figure 2).** Neuroinflammation is also a characteristic feature of AD pathogenesis linked to cognitive decline (21–23). MS cholinergic signaling controls hippocampal neuroinflammation and improves cognition in mice (24). We found age-dependent increases in the number of ionized calcium-binding adaptor protein-1 (IBA1)-positive microglia and morphological changes, indicative of neuroinflammation in female 5xFAD mice (**Supplementary Figure 3).** Collectively, these results demonstrate memory deficits in 5xFAD mice at 6 and 10 months of age, alongside reduced cholinergic neuronal body density in the MS and neuroinflammatory alterations in the hippocampus.

**Figure 1.**
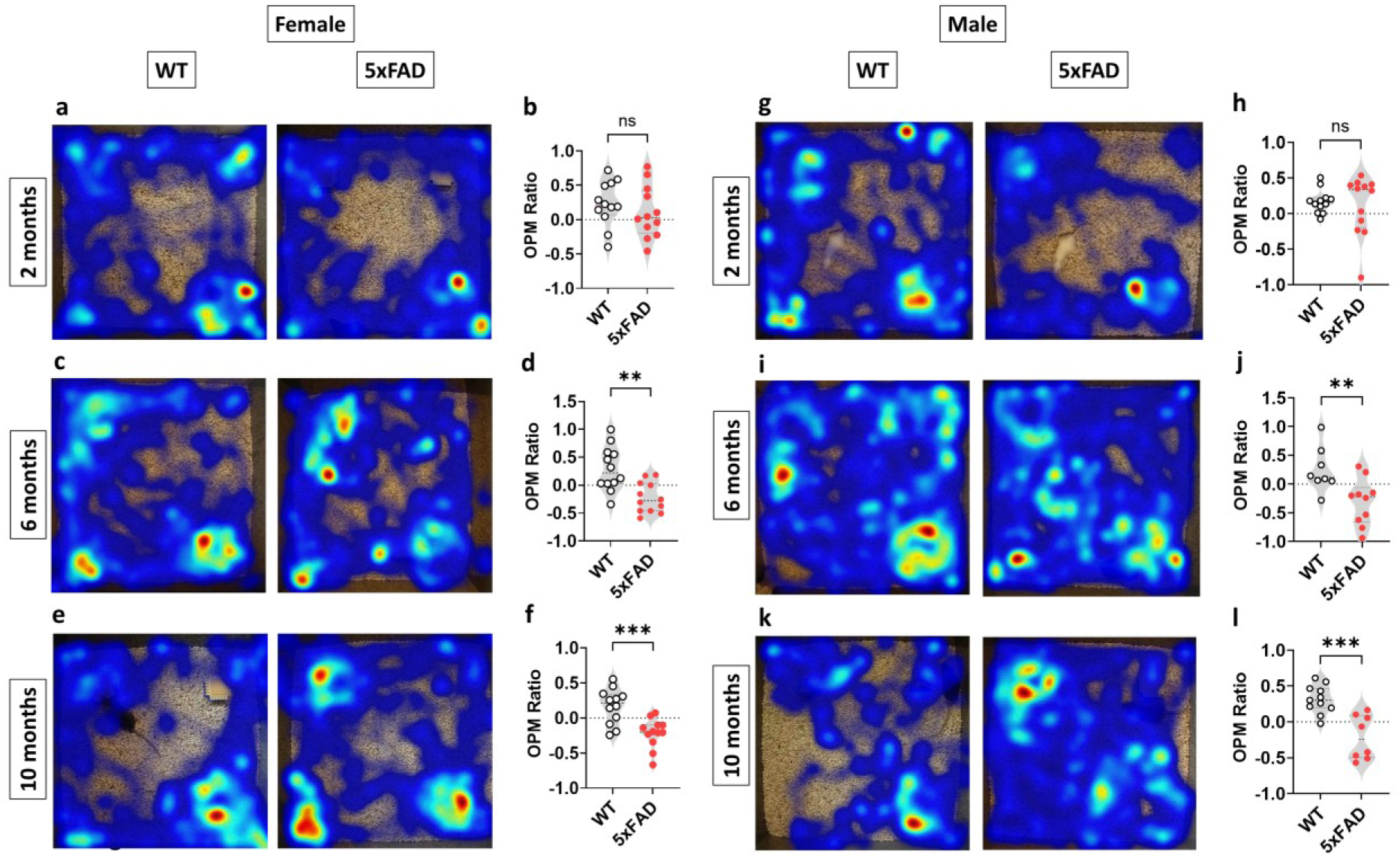
Age-dependent behavioral deficiencies in 5xFAD mice. OPM test is performed with 2-, 6-, and 10-month-old female and male WT and 5xFAD mice. (**a, b**) Representative occupancy heatmaps of female WT and 5xFAD mice during an OPM test and its analysis showing no difference in the OPM ratio between 2-month-old WT and 5xFAD mice (n=12, 12; Unpaired t test). (**c, d**) Representative occupancy heatmaps of 6-month-old female WT and 5xFAD mice during an OPM test and its analysis showing a significantly reduced OPM ratio in 5xFAD mice (n = 12,12; **p = 0.0011, Unpaired t test). (**e, f**) Representative occupancy heatmaps of 10-month-old female WT and 5xFAD mice during an OPM test and its analysis similarly show a significantly reduced OPM ratio in 5xFAD mice (n = 12,12; ***p = 0.0005, Unpaired t test). (**g, h**) Representative occupancy heatmaps of male WT and 5xFAD mice during an OPM test and its analysis showing no difference in the OPM ratio between 2-month-old WT and 5xFAD mice (n = 12,12; Unpaired t test). (**I, j**) Representative occupancy heatmaps of 6-month-old male WT and 5xFAD mice during an OPM test and its analysis showing a significantly reduced OPM ratio in 5xFAD mice (n = 8,10; **p = 0.009, Unpaired t test). (**k, l**) Representative occupancy heatmaps of 10-month-old male WT and 5xFAD mice during an OPM test and its analysis similarly show a significantly reduced OPM ratio in 5xFAD mice (n = 11,8; ***p = 0.0003, Unpaired t test). Data are presented as violin plots with individual mouse data points, and dotted horizontal lines indicating the median and interquartile range.

### Cholinergic neuron density in the dorsal motor nucleus of the vagus is reduced in an age-dependent manner in 5xFAD mice

Next, we localized the DMN on coronal sections using ChAT immunostaining and examined the DMN cholinergic neurons in female and male 5xFAD mice and control mice at 2, 6, and 10 months of age. In females at 2 months, there were no significant differences in the percentage of ChAT-positive neurons in the DMN between the 5xFAD and control mice **(Figure 2 a,b)**. However, at 6 months, the percentage of DMN ChAT-positive neurons was significantly reduced in female 5xFAD mice, compared with age-matched control mice (**Figure 2 c,d).** A decrease in the percentage of DMN ChAT-positive cholinergic neurons in female 5xFAD mice compared with controls was also observed at 10 months (**Figure 2 e,f).** Similar alterations were observed in male mice. As shown in **Figure 2 g,h,** no difference in the DMN ChAT-positive neurons was found at 2 months between 5xFAD and control mice. At 6 months, the percentage of DMN ChAT-positive neurons was significantly lower in the 5xFAD mice than in the age-matched control mice (**Figure 2 i,j**). At 10 months, these differences in ChAT-positive neuron percentage persisted; a significant decrease was observed in the DMN of 5xFAD mice compared with control mice (**Figure 2 k,l).** These results demonstrate significant reductions in cholinergic neuron density in the DMN of female and male 5xFAD mice at 6 and 10 months.

**Figure 2.**
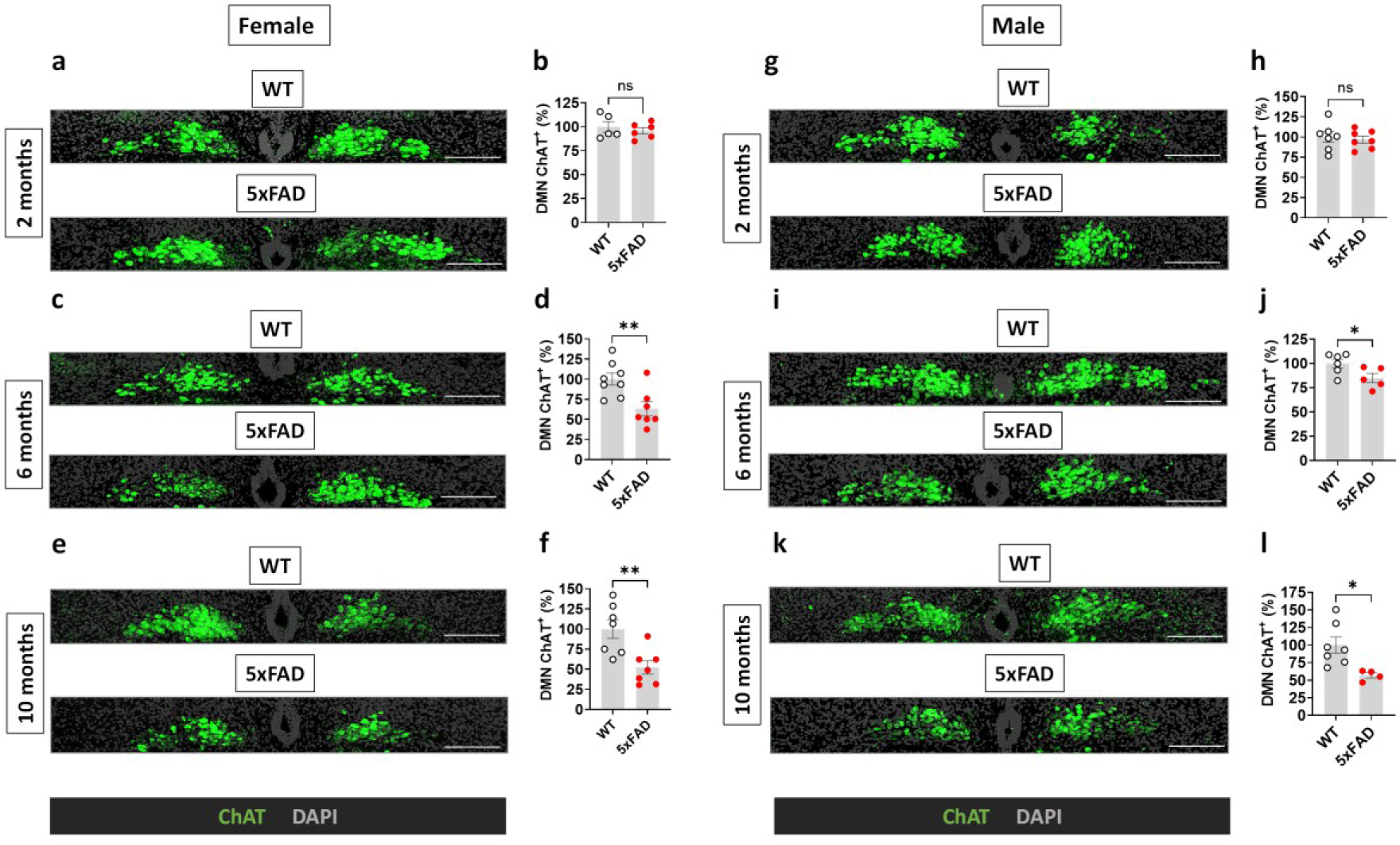
Age-dependent decrease of cholinergic (ChAT - positive) neuron density in the DMN of female and male 5xFAD mice. Cholinergic neurons in the DMN of 2-, 6-, and 10-month-old female and male WT and 5xFAD mice are identified by ChAT immunolabeling and quantified. (**a, b**) Representative images and quantitation showing no statistical difference in the percentage of ChAT-positive neurons in the DMN between 2-month-old WT and 5xFAD female mice (n = 5,6; Unpaired t test). Scale bar = 200 μm. (**c, d**) Representative images and quantitation demonstrating a significant decrease in ChAT-positive signal within the DMN of 5xFAD female mice at 6 months of age (n = 8,7; **p = 0.0066, Unpaired t test). (**e, f**) Representative images and quantitation showing a significant decrease in ChAT-positive neurons in the DMN of 5xFAD female mice at 10 months of age (n = 7,7; **p = 0.0057, Unpaired t test). (**g, h**) Representative images and quantitation showing no statistical difference in the percentage of ChAT-positive neurons in the DMN between 2-month-old WT and 5xFAD male mice (n = 7,7; Unpaired t test). (**I, j**) Representative images and quantitation demonstrating a significant decrease in ChAT-positive signal within the DMN of 5xFAD male mice at 6 months of age (n = 6,5; *p = 0.0439, Unpaired t test). (**k, l**) Representative images and quantitation showing a significant decrease in ChAT-positive neurons in the DMN of 5xFAD male mice at 10 months of age (n = 7, 4; *p = 0.0219, Unpaired t test). Each individual point represents the average of at least four, left and right combined technical replicates from an individual mouse. Data are presented as means ± SEM.

### Dorsal motor nucleus vagal control of the heart rate is impaired in an age-dependent manner in 5xFAD mice

Our observation of a significant reduction in the percentage of DMN cholinergic neurons over the course of the disease suggested impaired DMN vagal regulation. An important function of DMN cholinergic neurons in the efferent vagus nerve is cardiac regulation, including heart rate (25–30). We recently demonstrated that electrical DMN stimulation (eDMNS) significantly suppresses heart rate in male C57BL/6 mice (12). Others recently reported similar effects in transgenic mice subjected to optogenetic DMN stimulation (31). Therefore, we evaluated whether the eDMNS effect on heart rate differed by age in female and male 5xFAD mice compared with WT mice. To accomplish this, a stimulating electrode was inserted into the DMN, and heart rate was recorded for 1 min before, during, and after eDMNS in female mice (**Figure 3 a**). The heart rate recordings are shown in **Supplementary Figure 4**. As shown in **Figure 3 b,c,** the left side eDMNS (1mA, 260 μsec, 20Hz) significantly suppressed the heart rate in 2-month-old female WT and in 5xFAD mice. This effect of eDMNS was also demonstrated in 6-month-old female WT mice and was somewhat diminished in 5xFAD mice (**Figure 3 d,e)**. At 10 months, the significant effect of eDMNS on the heart rate in the WT female mice was significantly less manifested in 5xFAD mice (**Figure 3 f),** as demonstrated by the reduced bradycardia in 5xFAD mice compared with WT mice (**Figure 3 g**). Using the same experimental design, we examined the effects of eDMN stimulation on heart rate in male WT and 5xFAD mice (**Figure 4 a**). The heart rate recordings are shown in **Supplementary Figure 5**. We observed significant effects of eDMNS on heart rate in male WT mice at 2 and 6 months. The magnitude of this eDMNS effect was not significantly altered in male 5xFAD mice in the same 2-month and 6-month age groups (**Figure 4 b,c,d,e)**. However, at 10 months, this effect was significantly less pronounced in 5xFAD mice (**Figure 4 f**), as indicated by reduced relative heart rate (**Figure 4 g)**. These results indicate that the DMN-evoked bradycardia is significantly reduced in both female and male 10-month-old 5xFAD mice, consistent with diminished DMN cholinergic output.

**Figure 3.**
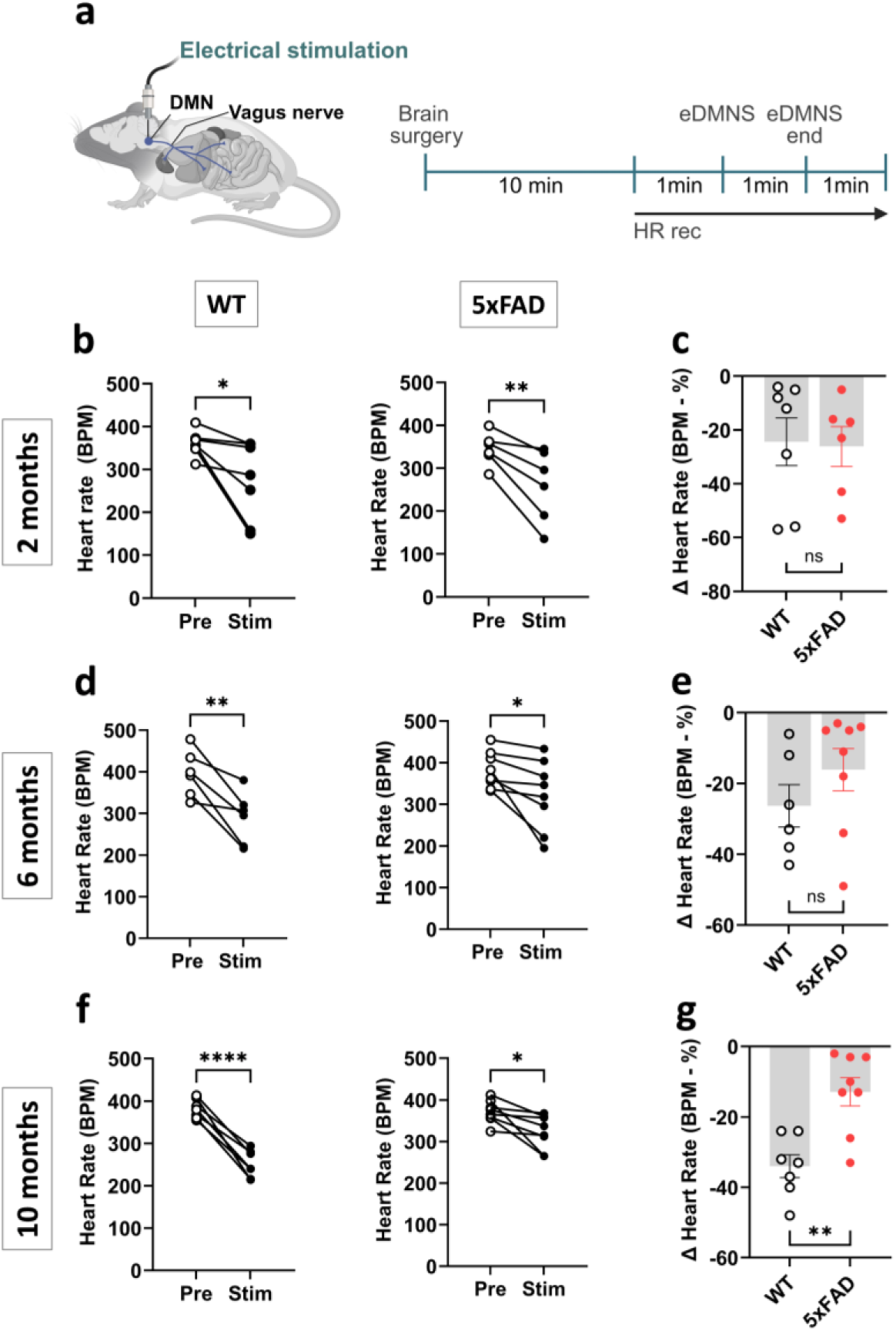
Age-dependent alterations in the suppressive effects of electrical stimulation of the dorsal motor nucleus of the vagus on the heart rate in female 5xFAD mice. (**a**) A schematic depiction of the experimental design. Anesthetized mice are placed on a stereotactic frame, brain surgical intervention is performed, and an electrode is inserted into the left DMN. Heart rate (HR) is recorded for 1 min prior to, during, and after electrical DMN stimulation (eDMNS) (Part of the schematic was created in BioRender). (**b**) eDMNS significantly suppresses the heart rate (HR) in 2-month-old female WT (n = 7, *p = 0.0361, Paired t test) and 5xFAD (n = 6, **p = 0.0098, Paired t test) mice. (**c**) The relative reduction in HR is comparable and non-significant (n = 7,6; Unpaired t test. (**d**) At 6 months of age, eDMNS significantly reduced HR in female WT (n = 6, **p = 0.0073, Paired t test) and 5xFAD (n = 8, *p = 0.0288, Paired t test) mice. (**e**) The relative reduction in HR is comparable and non-significant (n = 6,8; Mann-Whitney test). (**f**) eDMNS significantly reduces the HR of 10-month-old female WT (n = 7, ****p < 0.0001, Paired t test) and 5xFAD (n = 8, *p = 0.016, Paired t test) mice. (**g**) The relative reduction in HR for 5xFAD mice is significantly decreased (n = 7,8; **p = 0.0015, Unpaired t test). Data are presented as averaged HR, followed by a comparison of the relative percent drop. Each point represents one mouse. Data are presented as means ± SEM.

**Figure 4.**
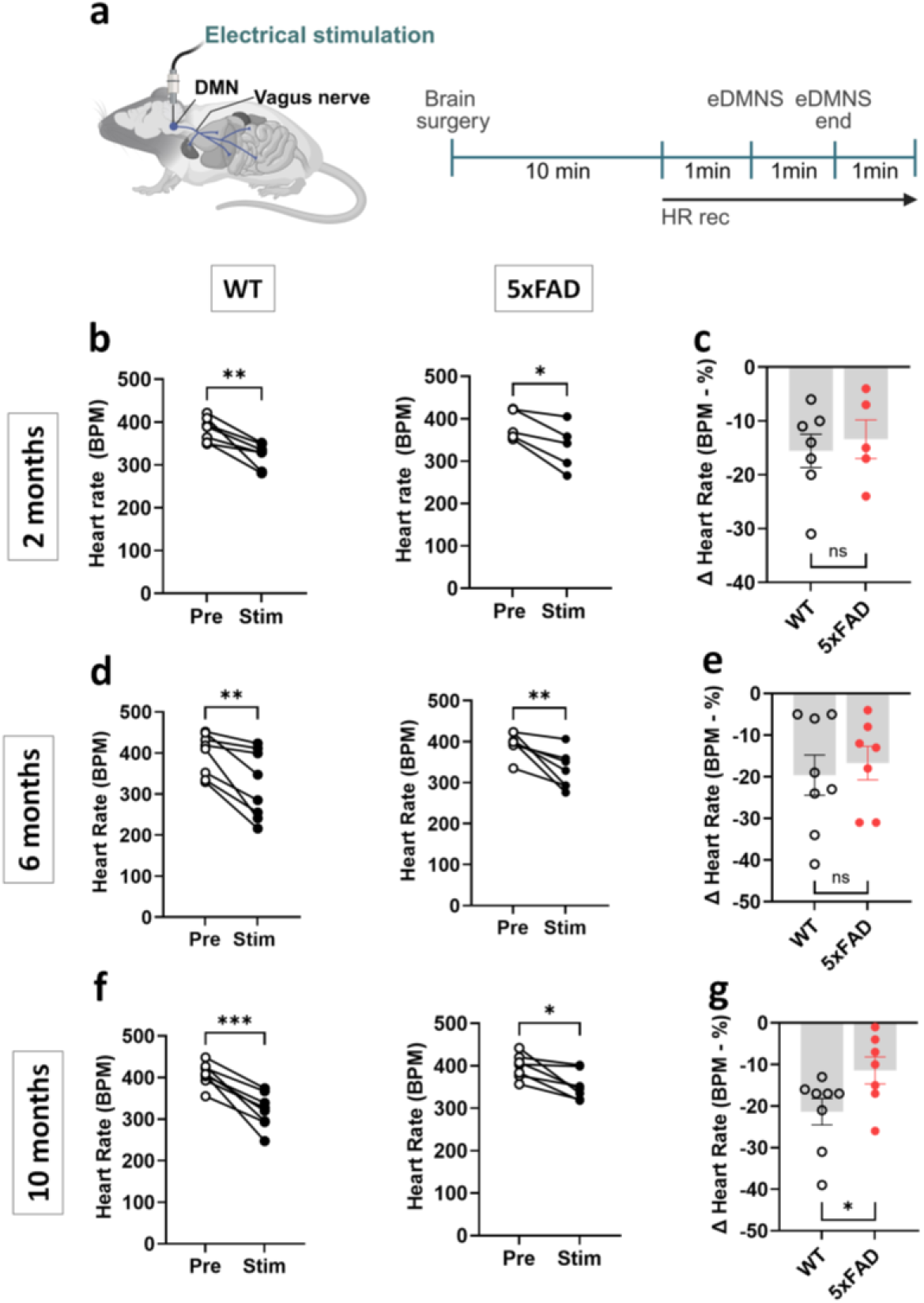
Age-dependent alterations in the suppressive effects of electrical stimulation of the DMN on the heart rate in male 5xFAD mice. **(a)** A schematic depiction of the experimental design (as described in Figure 3). (**b**) Electrical DMN stimulation (eDMNS) significantly suppresses the heart rate (HR) in 2-month-old male WT (n = 7; **p = 0.0034, Paired t test) and 5xFAD (n = 5; *p = 0.0165, Paired t test) mice. (**c**) The relative reduction in HR is comparable and non-significant (n = 7,5; Unpaired t test). (**d**) At 6 months of age, eDMNS significantly reduced HR in female WT (n = 8; **p = 0.0049, Paired t test) and 5xFAD (n = 7; **p = 0.0080, Paired t test) mice. (**e**) The relative reduction in HR is comparable and non-significant (n = 8,7; Unpaired t test). (**f**) eDMNS significantly reduces the HR of 10-month-old male WT (n = 8, ***p = 0.0003, Paired t test) and 5xFAD (n = 7, *p = 0.0144, Paired t test) mice. (**g**) The relative reduction in HR for 5xFAD mice is significantly decreased compared with WT mice (n = 8,7; *p = 0.042, Mann-Whitney test). Data are presented as averaged HR, followed by a comparison of the relative percent drop. Each point represents one mouse. Data are presented as means ± SEM.

### Dorsal motor nucleus vagal anti-inflammatory regulation is impaired in an age-dependent manner in 5xFAD mice

We recently showed that eDMNS reduced serum levels of the prototypical pro-inflammatory cytokine TNF in C57BL/6 mice during endotoxemia induced by lipopolysaccharide (LPS) i.p. administration (12). We also demonstrated that this effect was mediated through vagus nerve signaling (12). This study, together with the previously reported anti-inflammatory effect of optogenetic stimulation of DMN cholinergic neurons, characterized the DMN as a key source of efferent vagus cholinergic neurons that control inflammation (11, 12). Here, we studied the effect of eDMNS on serum TNF levels in 5xFAD mice and control mice subjected to endotoxemia across the three age groups. We previously demonstrated the anti-inflammatory effect of eDMNS (50 μA, 30Hz, 260 μsec) for 5 mins in male C57BL/6 mice only and did not perform experiments with female mice (12). Therefore, we first examined the effect of the same eDMNS regimen in female C57BL/6 mice (12). Using the same experimental procedure as before, in which left-sided eDMNS was performed prior to endotoxin administration, we showed that 5 min of eDMNS, compared with sham stimulation, significantly suppressed serum TNF levels in female mice during endotoxemia (**Supplementary Figure 6**). Then, we subjected female WT and 5xFAD mice aged 2, 6, and 10 months to sham stimulation or eDMNS prior to administering endotoxin (**Figure 5a**). As shown in **Figure 5 b,** eDMNS, compared with sham stimulation, significantly suppressed serum TNF levels in both 2-month-old WT and 5xFAD female mice. At 6 months, while eDMNS significantly lowered serum TNF levels in female WT mice, eDMNS in 5xFAD mice did not significantly alter the serum levels of this pro-inflammatory cytokine (**Figure 5 c**). Similarly, while eDMNS in 10-month-old female WT (control) mice significantly reduced serum TNF compared with sham stimulation, no significant alterations in serum TNF were observed when eDMNS was performed in 5xFAD mice during endotoxemia (**Figure 5 d).** In male WT and 5xFAD mice at 2 months of age, eDMNS (vs sham stimulation) significantly suppressed serum TNF levels during endotoxemia (**Figure 5 e**). At 6 months, eDMNS also significantly reduced serum TNF in both groups **(Figure 5 f).** However, at 10 months, although eDMNS significantly reduced serum TNF in WT mice, it failed to significantly alter serum TNF in 5xFAD male mice during endotoxemia (**Figure 5 g).** These results show that the eDMNS anti-inflammatory efficacy is significantly diminished at 6 and 10 months in female 5xFAD mice and at 10 months in male 5xFAD mice.

**Figure 5.**
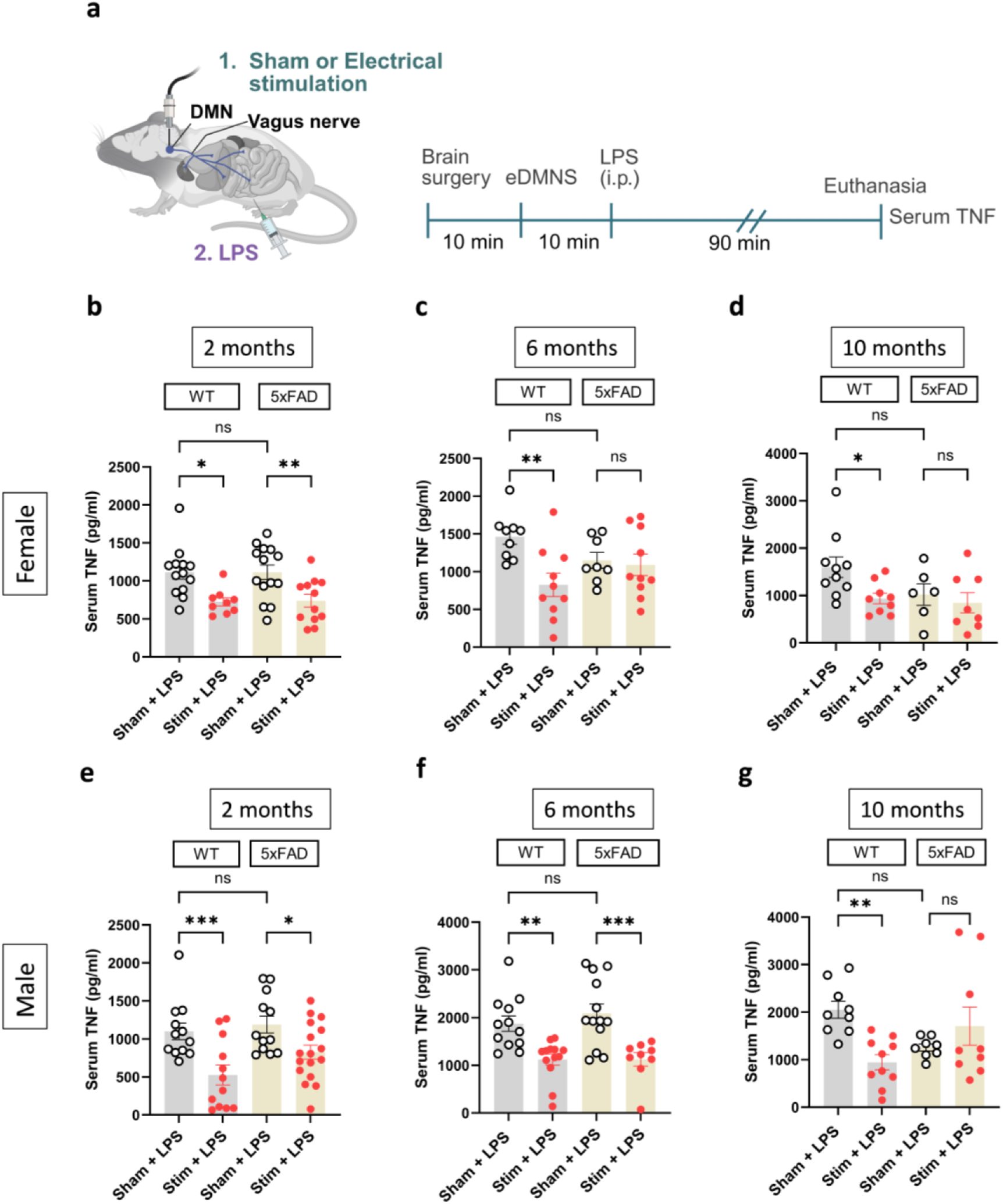
The suppressive effect of electrical dorsal motor nucleus of the vagus stimulation (eDMNS) on serum TNF levels in female and male WT mice is age-dependently diminished. **(a)** A schematic depiction of the experimental design. Anesthetized mice are placed on a stereotactic frame, and a brain surgical intervention is performed to insert an electrode into the left DMN. Sham stimulation or eDMNS is performed for 5 mins and LPS (0.5 mg/kg, i.p.) is administered to WT and 5xFAD mice. (Part of the schematic was created in BioRender) (**b**) At 2 months of age, eDMNS significantly reduces the LPS-induced serum TNF levels in both WT (n = 13,9; *p = 0.0104) and 5xFAD female mice (n=14,12; **p = 0.0062). (**c**) eDMNS in 6-month-old female mice significantly decreases serum TNF levels in WT mice (n = 9,10; **p = 0.003), but not in 5xFAD mice (n = 8,10). (**d**) Similarly, eDMNS decreases TNF in 10-month-old female WT mice (n = 11,9; *p = 0.0323) but not in 5xFAD mice (n = 6,8). (**e**) In male mice at 2 months of age, eDMNS suppresses serum TNF in WT (n = 12,12; ***p = 0.0007) and 5xFAD (n = 12,17; *p = 0.0377) mice. (**f**) eDMNS decreases serum TNF levels in 6-month-old WT (n = 12,13; **p = 0.0029) and 5xFAD (n = 13,9; ***p = 0.0005) male mice. (**g**) eDMNS in 10-month-old significantly reduces serum TNF levels in WT (n = 9,10; **p = 0.0041), but not in 5xFAD male mice (n = 8,7). Data are presented as individual mouse data points with means ± SEM. Data were analyzed using mixed effects two-way ANOVA with Sidak’s multiple comparisons.

## Discussion

Here, in mice exhibiting characteristic features of AD, including memory decline, basal forebrain cholinergic neurodegeneration, and hippocampal neuroinflammatory changes, we reveal previously unrecognized brainstem DMN cholinergic deficits, including reduced DMN cholinergic neuron density and weakened functional regulatory effects on heart rate and inflammation.

Both women and men suffer from AD, and there are sex dependent differences in terms of AD incidence, prevalence, and severity, with women being more affected (32–34). These findings necessitate research in both sexes. Accordingly, we carried out experiments with female and male 5xFAD mice, a widely utilized and well-characterized model of AD (37), and age-matched control mice. Importantly, in addition to age-dependent amyloid beta (Aβ) protein accumulation (amyloid plaques), a core feature of the 5xFAD model, other key AD pathologies, including cognitive deterioration, basal forebrain cholinergic neurodegeneration, and neuroinflammation, have been reported (35–39).

We first establish the timing of important AD features to provide a necessary time frame for studying DMN in 5xFAD mice. Memory decline linked to basal forebrain cholinergic neurodegeneration is a key clinical feature of AD, and there is a renewed interest in the cholinergic hypothesis of AD (2, 40, 41). Our results demonstrate memory (OPM) impairment in 5xFAD mice compared with controls at 6 and 10 months of age. These results are consistent with previously reported deterioration in object location memory in 5xFAD mice at 8 months, when mixed cohorts of female and male 4-month-old and 8-month-old mice were studied (17). MS cholinergic neurons innervate the hippocampus and regulate memory (20, 35, 42–44) and hippocampal microglial activation and neuroinflammation (24, 44). Our results show a significant cholinergic neuronal loss in the MS at 6 months and at 10 months in both female and male 5xFAD mice, in line with previous studies (35). We also show microglial changes, indicative of neuroinflammation, in the hippocampus of 6- and 10-month-old female 5xFAD mice. These observations corroborate previously reported hippocampal neuroinflammation, including an increase in the number of IBA1-positive microglia in male 5xFAD mice (39).

The anatomical and functional integrity of the brainstem DMN in AD has remained largely uncharacterized. Our results demonstrate age-dependent deterioration in brainstem DMN cholinergic neurons. While no differences are observed in 2-month-old female and male mice, the ChAT-positive neuronal signal is significantly decreased within the DMN at 6 months in 5xFAD mice of both sexes compared with controls. The decrease in cholinergic neurons is also significant at 10 months in both female and male mice. As the DMN cholinergic neurons are a major source of efferent vagus nerve fibers, these decreases should be associated with altered regulatory signaling through the vagus nerve. We show that eDMNS exerts weaker suppressive effects on heart rate in both female and male 10-month-old 5xFAD mice, indicating reduced DMN-mediated chronotropic control. At 6 months, eDMNS effects on heart rate are not significantly altered compared with those in control mice, indicating that even fewer DMN cholinergic neurons are sufficient to generate a regulatory effect of a similar magnitude. Another important function of DMN cholinergic neurons signaling via the vagus nerve is controlling pro-inflammatory cytokine responses, which we recently demonstrated using eDMNS in mice with systemic inflammation induced by LPS administration (12). Here, we show this eDMNS immunoregulatory effect (vs sham stimulation) on serum TNF levels in 2-month-old 5xFAD mice and WT mice of both sexes during endotoxemia. However, our results reveal that eDMNS fails to significantly alter serum TNF in 6-month-old female 5xFAD mice (vs sham stimulation), whereas this effect is retained in male 5xFAD mice of the same age. eDMNS fails to significantly suppress serum TNF in both female and male 10-month-old 5xFAD mice. These observations indicate an earlier impairment, at 6 months, of the anti-inflammatory function of DMN cholinergic neurons in female 5xFAD mice. Collectively, the diminished eDMNS suppressive effects on heart rate and TNF in 5xFAD mice support the biological relevance of the observed reductions in cholinergic (ChAT) signal in the DMN of female and male 5xFAD mice.

Numerous studies have documented autonomic dysfunction, cardiac imbalances, immune dysregulation and inflammation, and other peripheral alterations in AD (45–49). These peripheral changes occur in parallel with brain pathology in AD (21, 45–47, 49, 50), and peripheral inflammation has been shown to exacerbate neuroinflammation and the progression of AD (49, 51, 52). Autonomic dysfunction associated with diminished parasympathetic activity and vagal regulatory output to the heart has been documented in patients with AD (48, 53–57) and mild cognitive impairment (58). Furthermore, impaired cognitive performance in AD patients was associated with significantly decreased vagal and increased sympathetic heart regulation (59). While the mechanisms underlying autonomic dysfunction in AD remain poorly understood, it has been associated with alterations in brain networks and generalized underactivity of the cholinergic system (48, 54, 58). Our results in 5xFAD mice suggest that deterioration of DMN cholinergic neurons and diminished regulatory functions are likely to substantially contribute to autonomic dysfunction and peripheral derangements during AD and perhaps even play a causative role. Further studies should also extend to the brainstem nucleus ambiguus, another source of cholinergic vagal preganglionic fibers implicated in heart regulation. Impaired DMN vagal circuits may also promote other peripheral derangements, including inflammation that exacerbates neuroinflammation, which is linked to cognitive impairment in AD. The need to assess and treat autonomic dysfunction in AD patients to mitigate the potential risks has been acknowledged (60). A line of research that began with the discovery that electrical stimulation of the vagus nerve decreases serum TNF levels during murine endotoxemia (61) has demonstrated the anti-inflammatory efficacy of this approach in preclinical settings for many disorders (62, 63). Consecutive successful clinical trials have shown the anti-inflammatory and disease-alleviating effects of bioelectronic device-generated vagus nerve stimulation in chronic disorders (64–68), leading to the recent FDA approval of this therapeutic modality for rheumatoid arthritis. Non-invasive vagus neuromodulation has also been successfully evaluated in chronic inflammatory and autoimmune diseases (65, 69, 70). Accordingly, electrical vagus nerve stimulation and other approaches that counteract autonomic dysfunction and augment vagus nerve anti-inflammatory and beneficial cardiac effects (10, 61, 65) could be evaluated in therapeutic strategies for AD.

## Conclusion

Our results demonstrate anatomical deterioration of the brainstem DMN cholinergic neurons and functional impairment of DMN cholinergic vagal regulation of cardiac function and inflammation in a mouse model of AD. The timing of these age-dependent DMN alterations coincides with memory impairment and other established AD features. Revealing previously unrecognized DMN cholinergic dysfunction substantially advances our understanding of AD pathophysiology. These findings, along with further work in preclinical and clinical settings, can guide the development of novel AD treatments.

## Materials and methods

### Animals

All experimental procedures were approved by the Institutional Animal Care and Use Committee (IACUC) (IACUC protocol #2024-0065) of the Feinstein Institutes for Medical Research, Northwell Health, Manhasset, NY, in accordance with the NIH guidelines. We used a 5xFAD (Tg6799) Aβ-based mouse model. These mice are bred on the hybrid B6SJL background and express five AD-linked mutations of human APP and PSEN1 transgenes, including the Swedish (K670N/M671L), Florida (I716V), and London (V717I) mutations in APP, as well as the M146L and L286V mutations in PSEN1. Breeding mouse pairs were obtained from Jackson Laboratory and were allowed to acclimatize for three weeks prior to single-mate breeding. Breeding was carried out using female WT mice and male 5xFAD mice. Litters were weaned at 3-4 weeks of age and genotyped by Transnetyx (Cordova, TN) at this point. All female and male mice, positive (labeled as AD (5xFAD)) and negative (labeled as WT) for AD mutations, were raised until the experimental time points of 2, 6, or 10 months. All mice were given access to food and water *ad libitum* and maintained on a 12 h light and dark schedule at 25 °C and fed a standard Purina rodent chow diet (Rodents - Feed and nutrition products | Purina (multipurina.ca)).

### Immunohistochemistry and analysis

Female and male, WT and 5xFAD mice were euthanized with CO2 at 2, 6 or 10 months of age, transcardially perfused with PBS followed by 4% paraformaldehyde (PFA). Whole brains were removed and post-fixed overnight at 4 degrees Celsius in 4% PFA, prior to storage in 30% sucrose. The Paxinos and Franklin mouse brain atlas was used to define our brain regions of interest (the MS: bregma 1.09 - 0.61 mm; the hippocampus: bregma -1.55 - -2.03 mm; and the DMN: bregma -7.31 - -7.83 mm): A vibratome was used to obtain a series of 50 µm sections in our defined regions. Sections of defined brain regions were processed for immunohistochemical staining for ChAT and IBA1, respectively, within the defined brain region. The bregma-defined regions for the medial septum and the dorsal motor nucleus of the vagus were co-stained for ChAT and DAPI, whereas the hippocampal sections were stained for IBA1 and DAPI. Sections were washed, blocked (1% BSA), and as defined, incubated with primary antibodies, goat anti-ChAT (EMD Millipore AB144P: 1:100) and rabbit anti-IBA1 (Wako chemicals 019-19741: 1:400) for four nights at 4 degrees Celsius. Subsequently, sections were washed and incubated, as required, with fluorescent secondary antibodies donkey anti-goat 555 (Thermo fisher Scientific A21432: 1:200) and donkey anti-rabbit 647 (Thermo Fisher Scientific A31573: 1:500) for two nights at 4 degrees Celsius. Sections were briefly incubated with DAPI (1:10000) and then wet-mounted with Fluoromount. Slides were imaged using a Zeiss LSM 880 confocal microscope: tile-scans were utilized to represent ChAT expression in the MS and the DMN at 20x, and z-stacks were utilized to represent IBA1 activity in the hippocampus (left and right) at 40x.

Quantification of ChAT expression within Paxinos and Franklin mouse brain atlas-defined regions of the MS and the DMN was performed using region-of-interest selection and percent area positive for the ChAT signal in ImageJ. Quantification of IBA1 was performed using two complementary methods: a direct cell-body count to determine the number of microglia within defined brain regions, and a ramification index to describe microglial activity. The count was performed by inputting the average cell body size to ImageJ and using the counting feature to determine the quantity of microglia present; this result was also confirmed via manual counting. Ramification index was measured as previously described (71, 72). Using ImageJ, individual microglia had their outline traced (area), followed by a rough circle encompassing all their processes (perimeter). Area was divided by perimeter, providing a ‘ramification index’ - the closer the value is to 1, the more active the individual microglia (as defined by a typical pro-inflammatory phenotype). All images and quantification were performed by a blinded observer and were generally confirmed by a second blinded observer; typically, three replicates per animal were assessed. Specifically in the context of microglia, the ramification index of typically three microglia was ascertained, averaged and then subsequently averaged to replicate sections.

### Object place memory task

To assess spatial memory in 2-, 6-, and 10-month-old female and male WT and 5xFAD mice, the object place memory (OPM) task was performed as previously described (15). The testing apparatus consisted of a grey chamber with a square base (40 cm on each side) and 60-cm-high walls built of polyvinyl chloride. The floor was covered with a layer of bedding. A dim orange light was located above the chamber for illumination, as was a video camera for behavioral tracking.

Prior to initiation of behavioral experiments, mice were gently handled by the experimenters for a total of 45 min per mouse over the course of 3 days. Mice were acclimated to the testing apparatus, one at a time, over the course of four 15-min sessions (60 min total) spread across two days. The next day, mice were subjected, one at a time, to the OPM task, which consisted of three phases interspersed with 10-min intervals in the home cage. In the first phase, mice were placed into the testing apparatus for 15 min. This was followed by a sample trial in which two identical objects were placed in adjacent quadrants of the chamber, and the mice were returned to the chamber for 5 min. Finally, in the choice trial, one of the objects was moved to a new location in a different quadrant (moved object), while the other object remained in the same place (stable object), and the mouse was again returned to the chamber for 5 min. EthoVision behavioral tracking software was used to quantify the amount of time that the nose of each mouse was in proximity (<1 cm) to the periphery of either object, defined as the exploration time for that object. An ‘OPM ratio’ was calculated as (M – S)/(M + S), where M represents the amount of time the mouse spent exploring the moved object and S represents the amount of time the mouse spent exploring the stable object. A higher OPM ratio indicates more time spent exploring the moved object relative to the stable object and is interpreted as reflecting intact spatial memory in an individual mouse.

### Electrical DMN stimulation and heart rate measurement

Electrical DMN stimulation was performed according to the protocol reported in our previous study (12). The left DMN was localized as described previously in distinct 2-month-, 6-month-, and 10-month-old female and male WT (control) and 5xFAD mice. The animals were anesthetized by inhaling 2.5% isoflurane. Once fully sedated, the mice were positioned in a stereotaxic frame (Kopf) fitted with a nozzle for continuous isoflurane delivery at 1-1.5%. A midline incision was made in the skin, and the underlying muscles were retracted to expose the dura mater. The head was tilted forward to provide a clear view of the fourth ventricle through the dura mater. A slit was made in the dura mater with a 23G needle, the cerebrospinal fluid was drained, and the obex point was exposed. A concentric bipolar electrode, secured to the stereotaxic frame, was then inserted into the obex. As previously reported (13), the electrode was precisely guided to the left DMN coordinates defined in the Paxinos and Franklin mouse brain atlas (0.25 mm lateral to the obex and 0.48 mm). Mice were connected to a ‘small animal physiological monitoring system’ (Harvard apparatus) to measure heart rate, expressed as beats per minute (BPM). A stable baseline was obtained for each animal, and BPM was then recorded for 1 min pre-stimulation, 1 min during stimulation (1 mA, 30 Hz, 260 μsec), and 1 min immediately post-stimulation. Distinct graphs represent the data. The first graph shows BPM pre-, during, and post-stimulation. The second graph compares average BPM pre- and during stimulation. The third graph compares changes between WT and 5xFAD mice, with the average BPM expressed as a relative percentage drop from the baseline BPM (Δ change %).

### Electrical DMN stimulation, endotoxemia, and cytokine analysis

Female and male WT (controls) and 5xFAD mice at 2, 6, and 10 months of age were subjected to sham stimulation or eDMNS as previously reported (12). Briefly, anesthetized (with 2.5% isoflurane) mice were positioned in a stereotaxic frame, and a brain surgical procedure was performed as described above (for heart rate analysis). After positioning a bipolar electrode in the left DMN, stimulation was performed (in mice) or not (in sham mice) using a multi-channel STG4000 series stimulator at 50 μA, 30 Hz, 260 μsec for 5 min. Subsequently, the electrode was withdrawn, the animal was sutured, removed from the stereotaxic frame and lipopolysaccharide (LPS, endotoxin, Sigma-Aldrich, Serotype O111:B4, Lot# L4310-100 mg) (0.5 mg/kg) was administered intraperitoneally. Mice were allowed to recover in a heated cage for 90 mins, euthanized, and blood was collected and processed for analysis. Serum was analyzed for cytokine levels using a standard TNF ELISA kit (88–7324: Invitrogen). TNF was expressed as picograms per milliliter of serum collected.

### Statistics

Statistical analyses were conducted using GraphPad Prism 10 (GraphPad Software, Inc., La Jolla, CA). Data were assessed for normality using a Shapiro-Wilk test. Differences between two groups (analyzing immunohistochemistry and behavioral data) were assessed using an unpaired two-tailed Student’s *t* test (for normally distributed data) or the Mann-Whitney test (when the data did not meet the assumptions of normality). BPM was expressed as averaged individual traces, as the average BPM for pre-stimulation and stimulation, and, finally, as the BPM drop during stimulation converted to a percentage (Δ change %). Bar graph data were similarly assessed for normality. A paired t-test was performed for normally distributed data. For cytokine (TNF) analysis, a two-way ANOVA (mixed-effects analysis) was performed with Sidak’s multiple comparisons test. Data are presented as means ± standard error of the mean. P values less than 0.05 were considered significant for all experiments. All specific statistical tests are also listed in the figure legends.

## Acknowledgments

The study is reported in accordance with ARRIVE guidelines. The authors declare that no artificial intelligence (AI) tools were used in the preparation, data analysis, or figure generation of this manuscript. Some elements in the figures were generated via BioRender.com.

## Authors’ contributions

VAP and AF conceived the project. VAP, AF, JS, PH, SPP, and SC designed experiments. AF, SPP, SC, JJS, PTH, TT, and AT performed experiments and helped collect and process samples. AF, SSP, SC, JS, PH, AT, EC, and VAP analyzed data. All authors, including CD, LG, JC, YA, MB, SAC, and KJT, provided experimental advice, discussed the results, and commented on the manuscript. VAP wrote the first draft, and AF, MB, EC, and PM provided comments. All authors reviewed the manuscript and provided additional comments.

## Availability of data and materials

All data are provided in this manuscript.

## Supplementary Figures

**Supplementary Figure 1.**
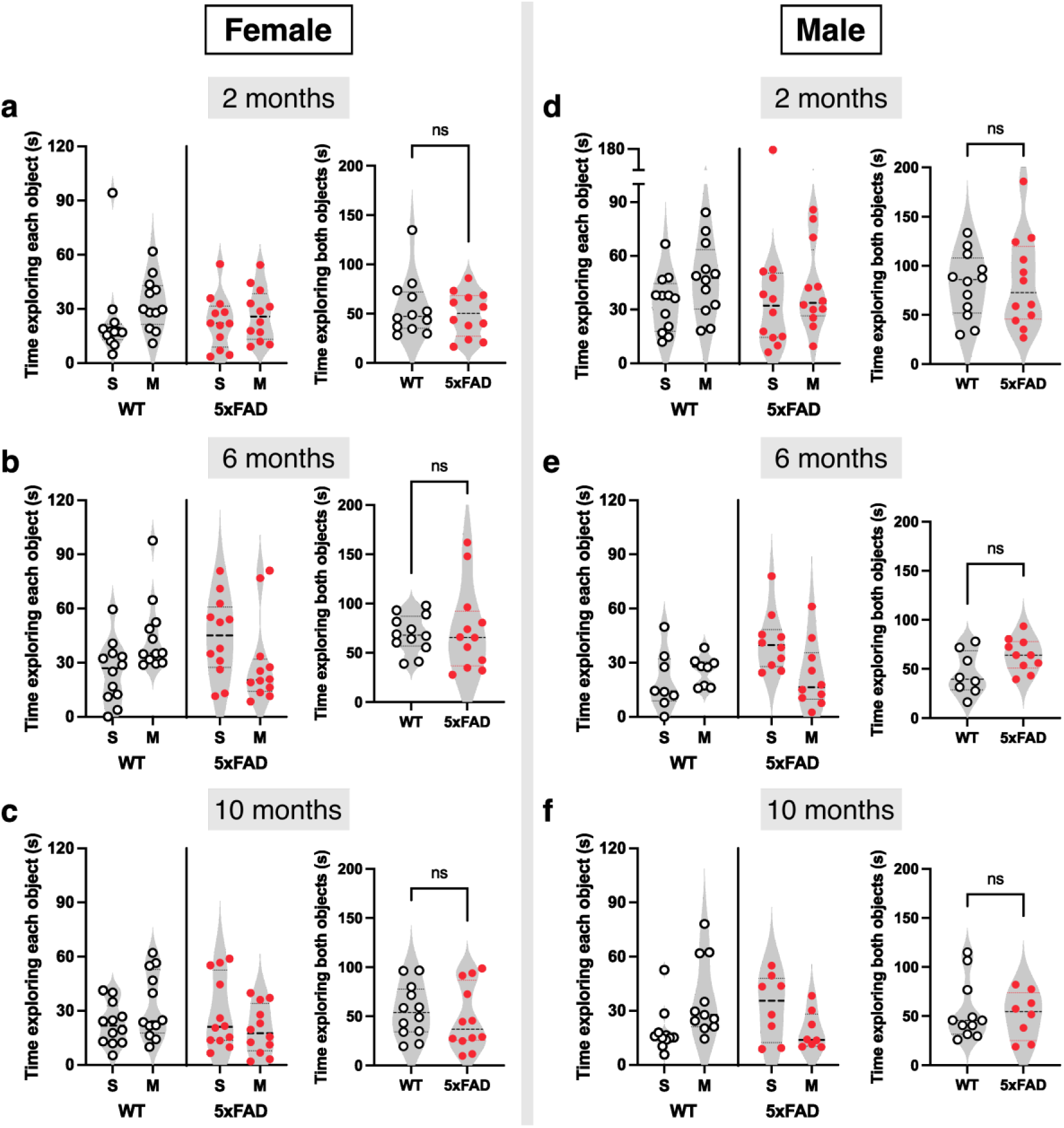
Comparable total object exploration times in WT and 5xFAD mice. No significant differences were detected between WT and 5xFAD mice in the total time spent exploring both objects during the choice trial in (**a**) 2-month-old females (n=12, 12; Unpaired t test), (**b**) 6-month-old females (n=12, 12; Unpaired t test), (**c**) 10-month-old females (n=12, 12; Unpaired t test), (**d**) 2-month-old males (n=12, 12; Unpaired t test), (**e**) 6-month-old males (n=8, 10; Unpaired t test), and (**f**) 10-month-old males (n=11, 8; Unpaired t test). For each age and sex group, the time spent exploring the stable object (S) and the moved object (M) is shown (*left*, *center*), as well as the total time spent exploring both objects (*right*). Data are presented as violin plots with individual mouse data points and dotted horizontal lines indicating the median and interquartile range.

**Supplementary Figure 2.**
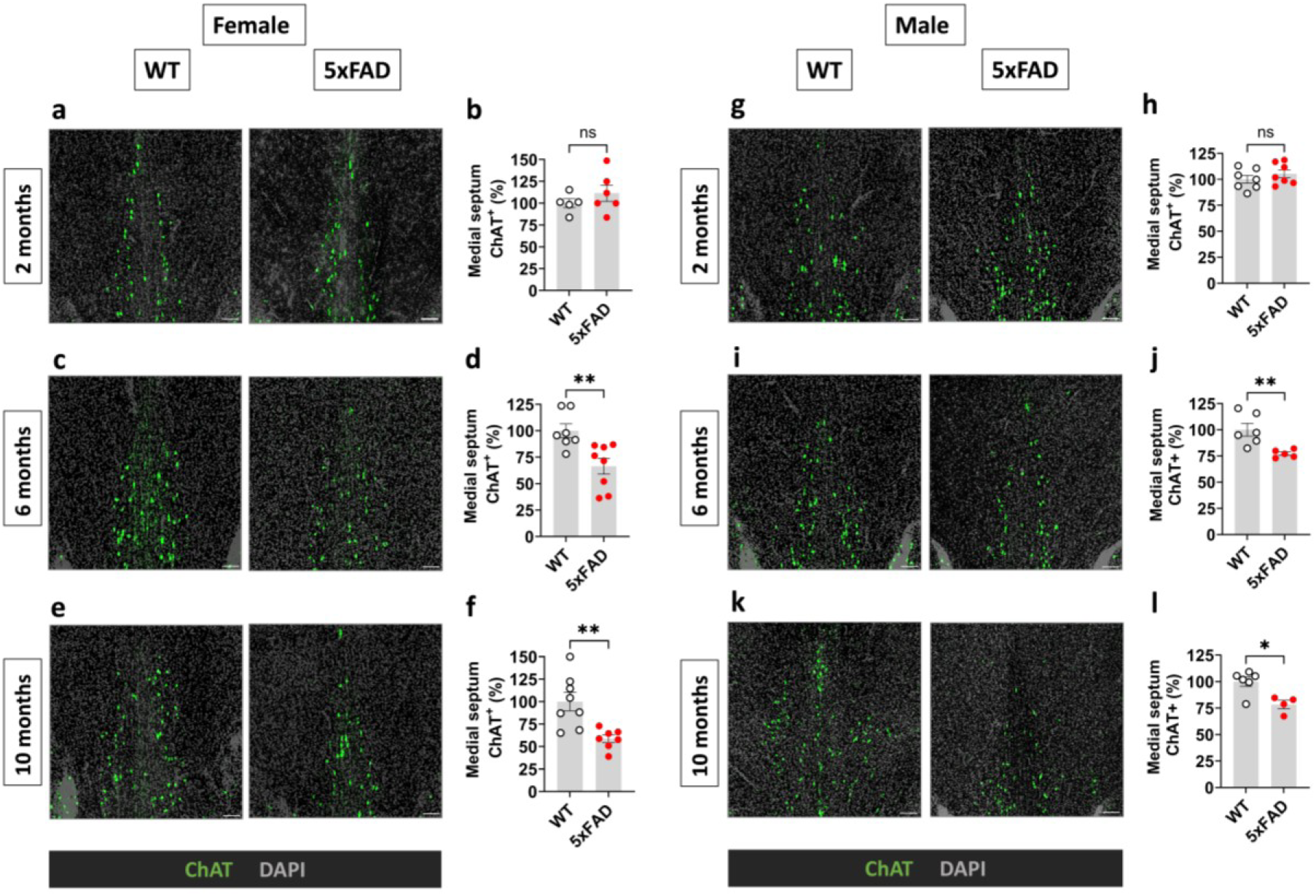
Age-dependent impaired integrity of cholinergic neurons in the MS of 5xFAD mice. Cholinergic neurons in the MS of 2-, 6-, and 10-month-old WT and 5xFAD female and male mice are identified by ChAT immunolabeling and quantified. (**a, b**) Representative images and quantitation showing no statistical difference in the percentage of ChAT-positive neurons in the MS between 2-month-old WT and 5xFAD female mice (n = 5,6; Unpaired t test). Scale bar = 200 μm. (**c,d**) Representative images and quantitation demonstrating a significant decrease in ChAT-positive signal within the MS of 5xFAD female mice at 6 months of age (n = 7,8; **p = 0.0056, Unpaired t test). (**e, f**) Representative images and quantitation showing a significant decrease in ChAT-positive neurons in the MS of 5xFAD female mice at 10 months of age (n = 8,7; **p = 0.0033, Unpaired t test). (**g, h**) Representative images and quantitation showing no statistical difference in the percentage of ChAT-positive neurons in the MS between 2-month-old WT and 5xFAD male mice (n = 7,7; Unpaired t test). (**I, j**) Representative images and quantitation demonstrating a significant decrease in ChAT-positive signal within the MS of 5xFAD male mice at 6 months of age (n = 6,5; **p = 0.0095, Unpaired t test). (**k, l**) Representative images and quantitation showing a significant decrease in ChAT-positive neurons in the MS of 5xFAD male mice at 10 months of age (n = 6,4; *p = 0.0381, Mann-Whitney test). Each point represents the average of 3 technical replicates from an individual mouse. Data are presented as means ± SEM.

**Supplementary Figure 3.**
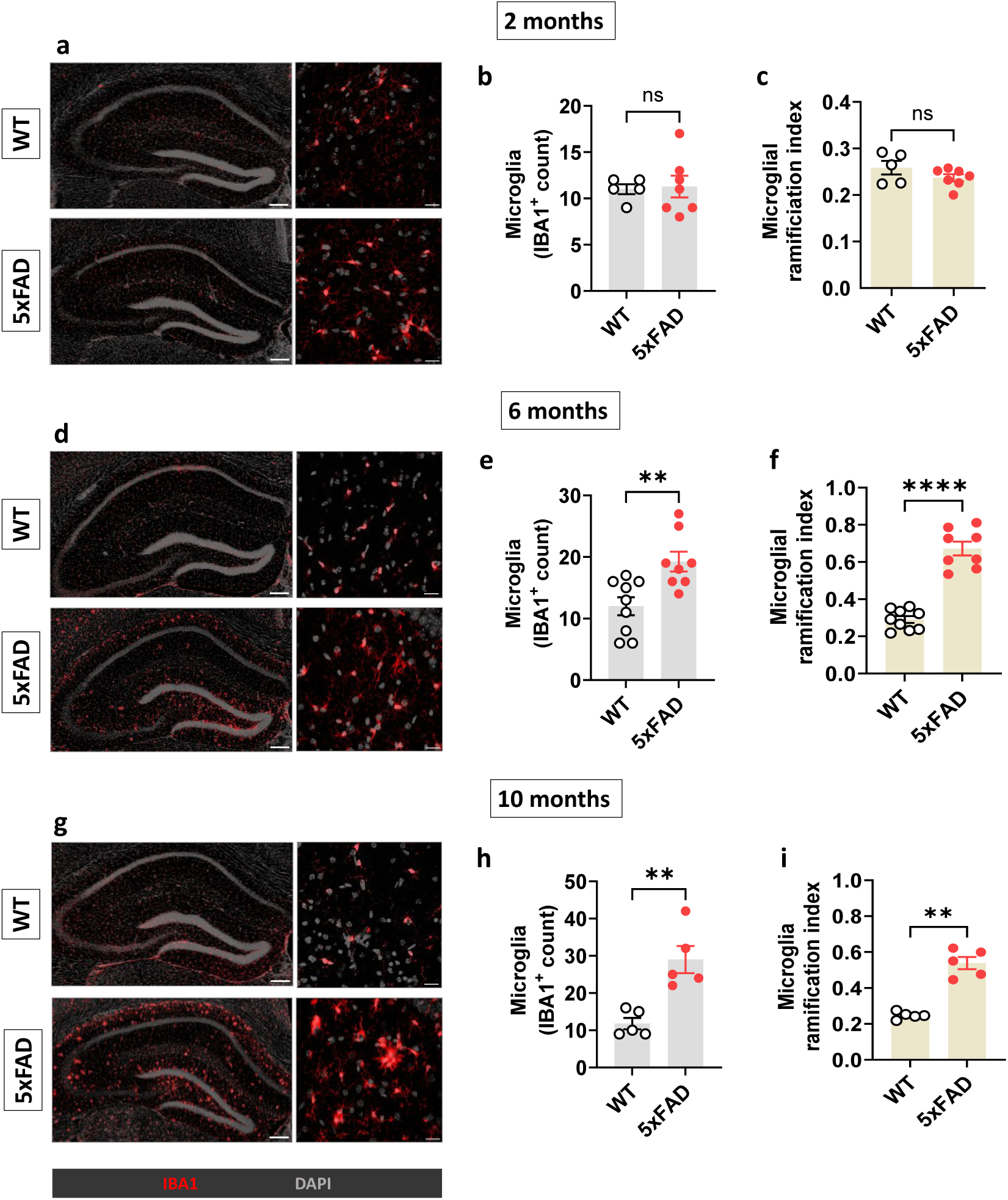
Age-dependent microglial alterations in the hippocampus of female 5xFAD mice. Microglia in the hippocampus of 2-, 6-, and 10-month-old female WT and 5xFAD mice are identified using IBA1 immunolabeling and quantified. (**a)** Representative images of hippocampal slices at 20x (left) and 40x (right) magnification for 2-month-old female WT (upper) and 5xFAD (lower) mice. Scale bar = 200 μm (tile scans) and 20 μm. At 2 months of age, there is no significant difference in the (**b**) number (n = 5,7; Unpaired t test) or (**c**) ramification of hippocampal microglia (n = 5,7; Unpaired t test) between WT and 5xFAD mice. (**d**) Representative images of hippocampal slices at 20x (left) and 40x (right) magnification for 6-month-old female WT (upper) and 5xFAD (lower) mice. There is a significantly greater (**e**) number of microglia (n = 9,8; **p = 0.0045, Unpaired t test), as well as (**f**) a higher ramification index (n = 9,8; ****p < 0.0001, Unpaired t test), in the hippocampus of 5xFAD mice compared with WT. (**g**) Representative images of hippocampal slices at 20x (left) and 40x (right) magnification for 10-month-old female WT (upper) and 5xFAD (lower) mice. There is a significantly greater (**h**) number of microglia (n = 5,5; **p = 0.0025, Unpaired t test), as well as (**I**) a higher ramification index (n = 5,5; **p = 0.0079, Mann-Whitney test), in the hippocampus of 5xFAD mice compared with WT. Data are presented as means ± SEM.

**Supplementary Figure 4.**
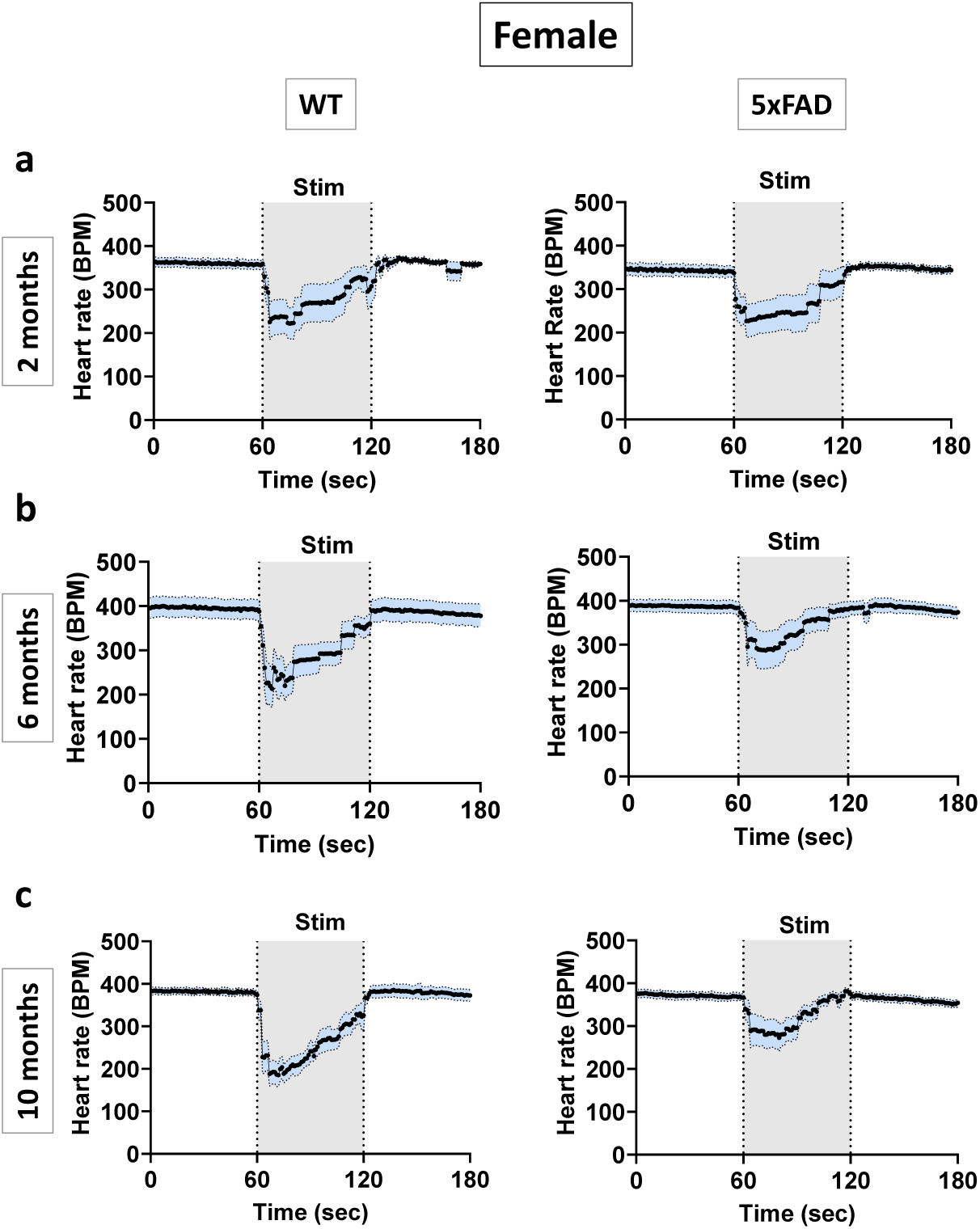
Heart rate recording before, during, and after electrical stimulation of the dorsal motor nucleus of the vagus in female wild-type (WT) and 5xFAD mice. **(a)** Heart rate recording for 2-month-old female mice. (**b**) Heart rate recording for 6-month-old female mice. (**c**) Heart rate recording for 10-month-old female mice. Data are represented as averaged HR traces from individual recordings.

**Supplementary Figure 5.**
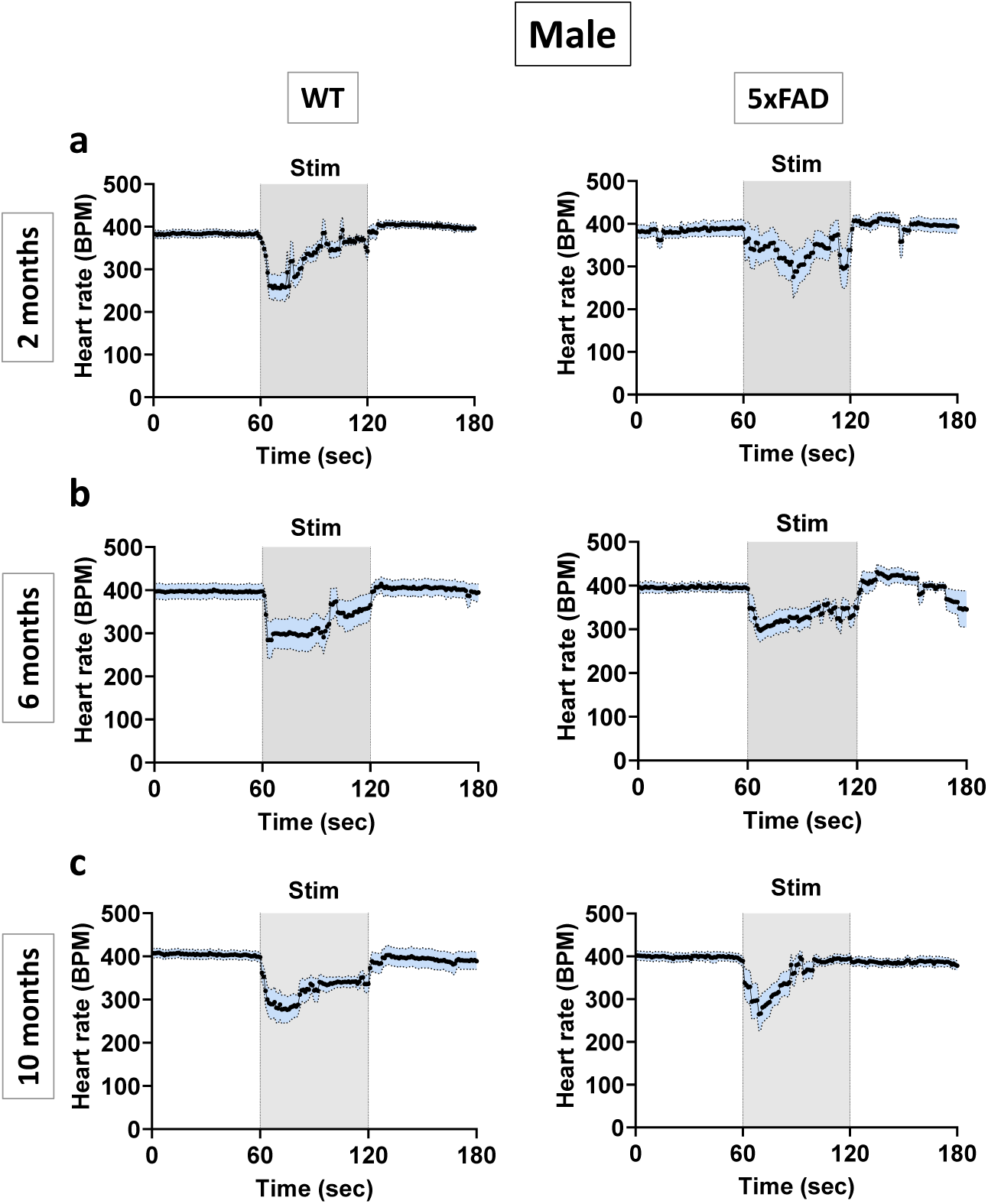
Heart rate recording before, during, and after electrical stimulation of the dorsal motor nucleus of the vagus in male wild-type (WT) and 5xFAD mice. **(a)** Heart rate recording for 2-month-old male mice. (**b**) Heart rate recording for 6-month-old male mice. (**c**) Heart rate recording for 10-month-old male mice. Data are represented as averaged HR traces from individual recordings.

**Supplementary Figure 6.**
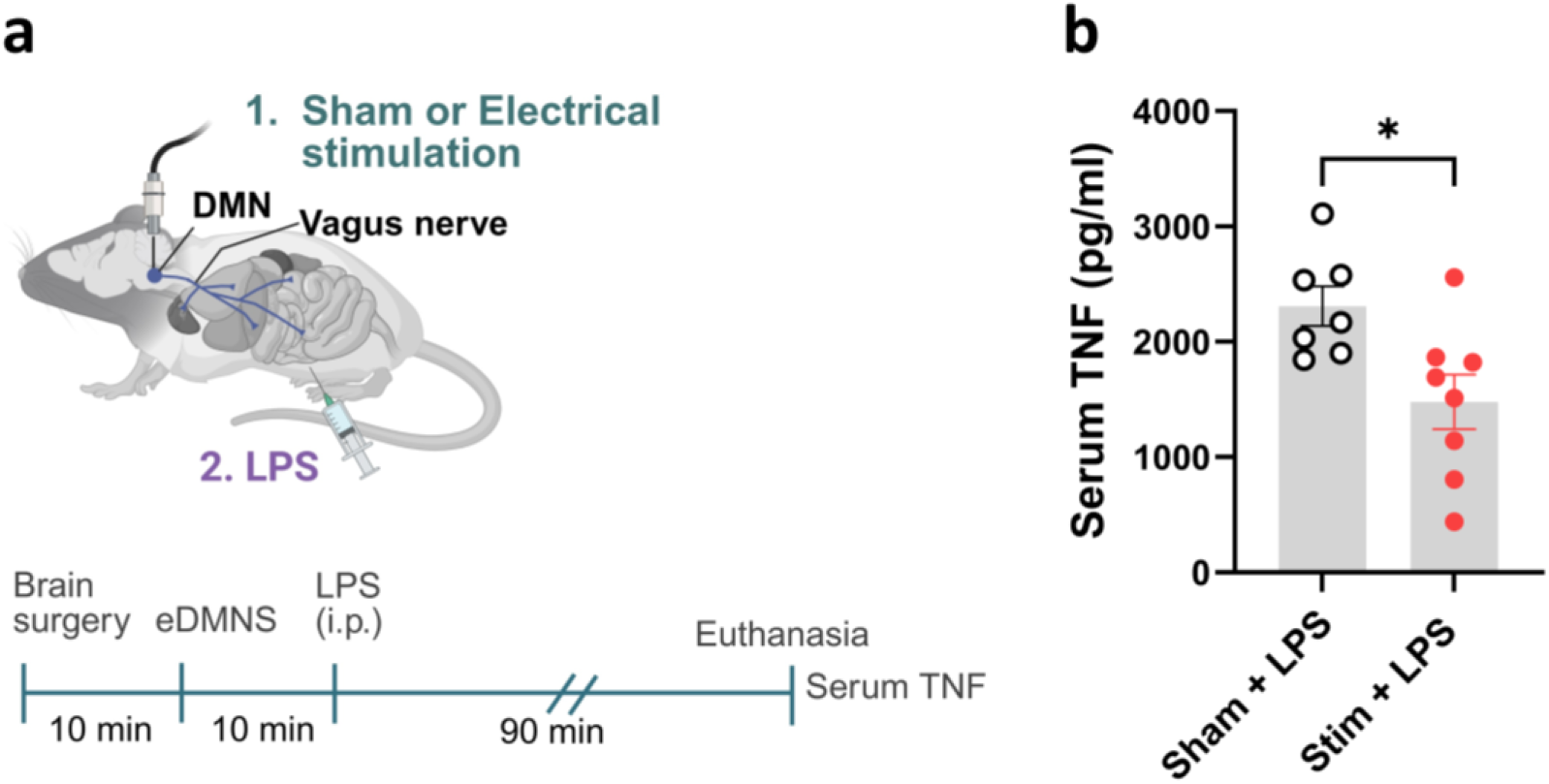
Electrical dorsal motor nucleus of the vagus stimulation (eDMNS) suppresses serum TNF levels in female C57BL/6 mice. (**a**) A schematic depiction of the experimental design. Anesthetized mice are placed in a stereotactic frame, and, through surgery, an electrode is inserted into the left DMN. Sham stimulation or eDMNS is performed for 5 mins, and LPS (0.5 mg/kg, i.p.) is administered. (**b**) eDMNS significantly decreases the expression of LPS-induced serum TNF levels in female mice (n = 7,8; *p = 0.016, Unpaired t test). Each point represents one mouse. Data are presented as means ± SEM.

## References

1. Brookmeyer R, Johnson E, Ziegler-Graham K, Arrighi HM. Forecasting the global burden of Alzheimer’s disease. Alzheimer’s & dementia : the journal of the Alzheimer’s Association. 2007;3(3):186–91.

2. Hampel H, Mesulam MM, Cuello AC, Farlow MR, Giacobini E, Grossberg GT, et al. The cholinergic system in the pathophysiology and treatment of Alzheimer’s disease. Brain. 2018;141(7):1917–33.

3. Livingston G, Sommerlad A, Orgeta V, Costafreda SG, Huntley J, Ames D, et al. Dementia prevention, intervention, and care. Lancet. 2017;390(10113):2673–734.

4. Bartus RT, Dean RL, 3rd, Beer B, Lippa AS. The cholinergic hypothesis of geriatric memory dysfunction. Science. 1982;217(4558):408–14.

5. Ballinger EC, Ananth M, Talmage DA, Role LW. Basal Forebrain Cholinergic Circuits and Signaling in Cognition and Cognitive Decline. Neuron. 2016;91(6):1199–218.

6. Birks J. Cholinesterase inhibitors for Alzheimer’s disease. The Cochrane database of systematic reviews. 2006(1):Cd005593.

7. Chang EH, Chavan SS, Pavlov VA. Cholinergic Control of Inflammation, Metabolic Dysfunction, and Cognitive Impairment in Obesity-Associated Disorders: Mechanisms and Novel Therapeutic Opportunities. Frontiers in neuroscience. 2019;13:263.

8. Pavlov VA, Tracey KJ. The vagus nerve and the inflammatory reflex--linking immunity and metabolism. Nature reviews Endocrinology. 2012;8(12):743–54.

9. Tracey KJ. The inflammatory reflex. Nature. 2002;420(6917):853–9.

10. Pavlov VA, Tracey KJ. Neural regulation of immunity: molecular mechanisms and clinical translation. Nature neuroscience. 2017;20(2):156–66.

11. Kressel AM, Tsaava T, Levine YA, Chang EH, Addorisio ME, Chang Q, et al. Identification of a brainstem locus that inhibits tumor necrosis factor. Proceedings of the National Academy of Sciences of the United States of America. 2020;117(47):29803–10.

12. Falvey A, Palandira SP, Chavan SS, Brines M, Dantzer R, Tracey KJ, et al. Electrical stimulation of the dorsal motor nucleus of the vagus in male mice can regulate inflammation without affecting the heart rate. Brain Behav Immun. 2024.

13. Greene JG. Causes and consequences of degeneration of the dorsal motor nucleus of the vagus nerve in Parkinson’s disease. Antioxid Redox Signal. 2014;21(4):649–67.

14. Zokaei N, Sillence A, Kienast A, Drew D, Plant O, Slavkova E, et al. Different patterns of short-term memory deficit in Alzheimer’s disease, Parkinson’s disease and subjective cognitive impairment. Cortex. 2020;132:41–50.

15. Faust TW, Robbiati S, Huerta TS, Huerta PT. Dynamic NMDAR-mediated properties of place cells during the object place memory task. Front Behav Neurosci. 2013;7:202.

16. Assini FL, Duzzioni M, Takahashi RN. Object location memory in mice: Pharmacological validation and further evidence of hippocampal CA1 participation. Behav Brain Res. 2009;204(1):206–11.

17. Zhang H, Chen L, Johnston KG, Crapser J, Green KN, Ha NM-L, et al. Degenerate mapping of environmental location presages deficits in object-location encoding and memory in the 5xFAD mouse model for Alzheimer’s disease. Neurobiol Dis. 2023;176:105939.

18. Murai T, Okuda S, Tanaka T, Ohta H. Characteristics of object location memory in mice: Behavioral and pharmacological studies. Physiol Behav. 2007;90(1):116–24.

19. Ikonen S, McMahan R, Gallagher M, Eichenbaum H, Tanila H. Cholinergic system regulation of spatial representation by the hippocampus. Hippocampus. 2002;12(3):386–97.

20. Mamad O, McNamara HM, Reilly RB, Tsanov M. Medial septum regulates the hippocampal spatial representation. Front Behav Neurosci. 2015;9:166.

21. Heneka MT, Carson MJ, El Khoury J, Landreth GE, Brosseron F, Feinstein DL, et al. Neuroinflammation in Alzheimer’s disease. The Lancet Neurology. 2015;14(4):388–405.

22. Calsolaro V, Edison P. Neuroinflammation in Alzheimer’s disease: Current evidence and future directions. Alzheimer’s & Dementia. 2016;12(6):719–32.

23. Leng F, Edison P. Neuroinflammation and microglial activation in Alzheimer disease: where do we go from here? Nature Reviews Neurology. 2021;17(3):157–72.

24. Yin L, Zhang J, Ma H, Zhang X, Fan Z, Yang Y, et al. Selective activation of cholinergic neurotransmission from the medial septal nucleus to hippocampal pyramidal neurones improves sepsis-induced cognitive deficits in mice. Br J Anaesth. 2023.

25. Machhada A, Ang R, Ackland GL, Ninkina N, Buchman VL, Lythgoe MF, et al. Control of ventricular excitability by neurons of the dorsal motor nucleus of the vagus nerve. Heart Rhythm. 2015;12(11):2285–93.

26. Gourine AV, Machhada A, Trapp S, Spyer KM. Cardiac vagal preganglionic neurones: An update. Autonomic Neuroscience. 2016;199:24–8.

27. Machhada A, Marina N, Korsak A, Stuckey DJ, Lythgoe MF, Gourine AV. Origins of the vagal drive controlling left ventricular contractility. J Physiol. 2016;594(14):4017–30.

28. Pickering A, Salo L, Hewinson J, Ambler M, Paton J, McAllen R. Does the dorsal motor nucleus of the vagus control cardiac chronotropism in the rat? Autonomic Neuroscience. 2011;163(1-2):120.

29. Geis GS, Wurster RD. Cardiac responses during stimulation of the dorsal motor nucleus and nucleus ambiguus in the cat. Circulation Research. 1980;46(5):606–11.

30. Jones JFX, Wang Y, Jordan D. Activity of C fibre cardiac vagal efferents in anaesthetized cats and rats. The Journal of Physiology. 1998;507(3):869–80.

31. Strain MM, Conley NJ, Kauffman LS, Espinoza L, Fedorchak S, Martinez PC, et al. Dorsal motor vagal neurons can elicit bradycardia and reduce anxiety-like behavior. iScience. 2024;27(3):109137.

32. Ferretti MT, Iulita MF, Cavedo E, Chiesa PA, Schumacher Dimech A, Santuccione Chadha A, et al. Sex differences in Alzheimer disease - the gateway to precision medicine. Nature reviews Neurology. 2018;14(8):457–69.

33. Zhu D, Montagne A, Zhao Z. Alzheimer’s pathogenic mechanisms and underlying sex difference. Cellular and molecular life sciences : CMLS. 2021;78(11):4907–20.

34. 2020 Alzheimer’s disease facts and figures. Alzheimer’s & Dementia. 2020;16(3):391–460.

35. Devi L, Ohno M. Phospho-eIF2α level is important for determining abilities of BACE1 reduction to rescue cholinergic neurodegeneration and memory defects in 5XFAD mice. PLoS One. 2010;5(9):e12974.

36. Yan H, Pang P, Chen W, Zhu H, Henok KA, Li H, et al. The Lesion Analysis of Cholinergic Neurons in 5XFAD Mouse Model in the Three-Dimensional Level of Whole Brain. Molecular neurobiology. 2018;55(5):4115–25.

37. Oakley H, Cole SL, Logan S, Maus E, Shao P, Craft J, et al. Intraneuronal beta-amyloid aggregates, neurodegeneration, and neuron loss in transgenic mice with five familial Alzheimer’s disease mutations: potential factors in amyloid plaque formation. The Journal of neuroscience : the official journal of the Society for Neuroscience. 2006;26(40):10129–40.

38. Oblak AL, Lin PB, Kotredes KP, Pandey RS, Garceau D, Williams HM, et al. Comprehensive Evaluation of the 5XFAD Mouse Model for Preclinical Testing Applications: A MODEL-AD Study. Frontiers in aging neuroscience. 2021;13:713726.

39. Mezö C, Dokalis N, Mossad O, Staszewski O, Neuber J, Yilmaz B, et al. Different effects of constitutive and induced microbiota modulation on microglia in a mouse model of Alzheimer’s disease. Acta Neuropathol Commun. 2020;8(1):119.

40. Berry AS, Harrison TM. New perspectives on the basal forebrain cholinergic system in Alzheimer’s disease. Neurosci Biobehav Rev. 2023;150:105192.

41. Metz CN, Pavlov VA. Treating disorders across the lifespan by modulating cholinergic signaling with galantamine. Journal of neurochemistry. 2021;158(6):1359–80.

42. Müller C, Remy S. Septo-hippocampal interaction. Cell Tissue Res. 2018;373(3):565–75.

43. Wang Y, Shen Y, Cai X, Yu J, Chen C, Tan B, et al. Deep brain stimulation in the medial septum attenuates temporal lobe epilepsy via entrainment of hippocampal theta rhythm. CNS Neuroscience & Therapeutics. 2021;27(5):577–86.

44. Dobryakova YV, Volobueva MN, Manolova AO, Medvedeva TM, Kvichansky AA, Gulyaeva NV, et al. Cholinergic Deficit Induced by Central Administration of 192IgG-Saporin Is Associated With Activation of Microglia and Cell Loss in the Dorsal Hippocampus of Rats. Frontiers in neuroscience. 2019;13:146.

45. Morris G, Berk M, Maes M, Puri BK. Could Alzheimer’s Disease Originate in the Periphery and If So How So? Molecular neurobiology. 2019;56(1):406–34.

46. Kowalski K, Mulak A. Brain-Gut-Microbiota Axis in Alzheimer’s Disease. Journal of neurogastroenterology and motility. 2019;25(1):48–60.

47. de Bruijn RF, Ikram MA. Cardiovascular risk factors and future risk of Alzheimer’s disease. BMC medicine. 2014;12:130.

48. Femminella GD, Rengo G, Komici K, Iacotucci P, Petraglia L, Pagano G, et al. Autonomic dysfunction in Alzheimer’s disease: tools for assessment and review of the literature. Journal of Alzheimer’s disease : JAD. 2014;42(2):369–77.

49. Yang J, Liang J, Hu N, He N, Liu B, Liu G, et al. The Gut Microbiota Modulates Neuroinflammation in Alzheimer’s Disease: Elucidating Crucial Factors and Mechanistic Underpinnings. CNS Neurosci Ther. 2024;30(10):e70091.

50. McManus RM, Heneka MT. Role of neuroinflammation in neurodegeneration: new insights. Alzheimer’s research & therapy. 2017;9(1):14.

51. Kitazawa M, Oddo S, Yamasaki TR, Green KN, LaFerla FM. Lipopolysaccharide-induced inflammation exacerbates tau pathology by a cyclin-dependent kinase 5-mediated pathway in a transgenic model of Alzheimer’s disease. J Neurosci. 2005;25(39):8843–53.

52. Liu Y, Zhang S, Li X, Liu E, Wang X, Zhou Q, et al. Peripheral inflammation promotes brain tau transmission via disrupting blood–brain barrier. Biosci Rep. 2020;40(2):BSR20193629.

53. Giubilei F, Strano S, Imbimbo BP, Tisei P, Calcagnini G, Lino S, et al. Cardiac autonomic dysfunction in patients with Alzheimer disease: possible pathogenetic mechanisms. Alzheimer Dis Assoc Disord. 1998;12(4):356–61.

54. Royall DR, Gao JH, Kellogg DL, Jr. Insular Alzheimer’s disease pathology as a cause of “age-related” autonomic dysfunction and mortality in the non-demented elderly. Med Hypotheses. 2006;67(4):747–58.

55. Toledo MA, Junqueira LF, Jr. Cardiac autonomic modulation and cognitive status in Alzheimer’s disease. Clin Auton Res. 2010;20(1):11–7.

56. Zulli R, Nicosia F, Borroni B, Agosti C, Prometti P, Donati P, et al. QT Dispersion and Heart Rate Variability Abnormalities in Alzheimer’s Disease and in Mild Cognitive Impairment. Journal of the American Geriatrics Society. 2005;53(12):2135–9.

57. Geng D, Wang Q, Zheng W, Yin Y, Xu P, Xu G. ECG- and HRV-Based Hybrid Architecture—Early Detection of Alzheimer’s Disease and Mild Cognitive Impairment. Applied Sciences. 2025;15(23):12555.

58. Collins O, Dillon S, Finucane C, Lawlor B, Kenny RA. Parasympathetic autonomic dysfunction is common in mild cognitive impairment. Neurobiol Aging. 2012;33(10):2324–33.

59. Nonogaki Z, Umegaki H, Makino T, Suzuki Y, Kuzuya M. Relationship between cardiac autonomic function and cognitive function in Alzheimer’s disease. Geriatrics & Gerontology International. 2017;17(1):92–8.

60. Papadopoulou M, Stefanou MI, Bakola E, Moschovos C, Athanasaki A, Tsigkaropoulou E, et al. Dysautonomia in Alzheimer’s Disease: A Systematic Review. Brain Sci. 2025;15(5).

61. Borovikova LV, Ivanova S, Zhang M, Yang H, Botchkina GI, Watkins LR, et al. Vagus nerve stimulation attenuates the systemic inflammatory response to endotoxin. Nature. 2000;405(6785):458–62.

62. Falvey A, Metz CN, Tracey KJ, Pavlov VA. Peripheral nerve stimulation and immunity: the expanding opportunities for providing mechanistic insight and therapeutic intervention. International Immunology. 2022;34(2):107–18.

63. Pavlov VA, Chavan SS, Tracey KJ. Molecular and Functional Neuroscience in Immunity. Annual review of immunology. 2018;36:783–812.

64. Tesser JRP, Crowley AR, Box EJ, June JP, Wickersham PB, Valenzuela GJ, et al. Vagus nerve-mediated neuroimmune modulation for rheumatoid arthritis: a pivotal randomized controlled trial. Nat Med. 2025.

65. Pavlov VA, Tracey KJ. Bioelectronic medicine: Preclinical insights and clinical advances. Neuron. 2022;110(21):3627–44.

66. Koopman FA, Chavan SS, Miljko S, Grazio S, Sokolovic S, Schuurman PR, et al. Vagus nerve stimulation inhibits cytokine production and attenuates disease severity in rheumatoid arthritis. Proceedings of the National Academy of Sciences of the United States of America. 2016;113(29):8284–9.

67. Bonaz B, Sinniger V, Hoffmann D, Clarencon D, Mathieu N, Dantzer C, et al. Chronic vagus nerve stimulation in Crohn’s disease: a 6-month follow-up pilot study. Neurogastroenterology and motility : the official journal of the European Gastrointestinal Motility Society. 2016;28(6):948–53.

68. D’Haens G, Eberhardson M, Cabrijan Z, Danese S, van den Berg R, Löwenberg M, et al. Neuroimmune modulation through vagus nerve stimulation reduces inflammatory activity in Crohn’s disease patients: a prospective open label study. Journal of Crohn’s and Colitis. 2023.

69. Aranow C, Atish-Fregoso Y, Lesser M, Mackay M, Anderson E, Chavan S, et al. Transcutaneous auricular vagus nerve stimulation reduces pain and fatigue in patients with systemic lupus erythematosus: a randomised, double-blind, sham-controlled pilot trial. Annals of the Rheumatic Diseases. 2021;80(2):203–8.

70. Tynan A, Brines M, Chavan SS. Control of inflammation using non-invasive neuromodulation: past, present and promise. Int Immunol. 2022;34(2):119–28.

71. Wittekindt M, Kaddatz H, Joost S, Staffeld A, Bitar Y, Kipp M, et al. Different Methods for Evaluating Microglial Activation Using Anti-Ionized Calcium-Binding Adaptor Protein-1 Immunohistochemistry in the Cuprizone Model. Cells. 2022;11(11):1723.

72. Falvey A, Palandira SP, Chaudhry S, Tynan A, Consolim-Colombo FM, Metz CN, et al. The cholinergic drug galantamine ameliorates acute and subacute peripheral and brain manifestations of acute respiratory distress syndrome in mice. Scientific Reports. 2025;15(1):33217.

